# Melatonin alleviates LPS-induced abnormal pregnancy through MTNR1B regulation of m6A

**DOI:** 10.1101/2023.04.19.537547

**Authors:** Shisu Zhao, Yanjun Dong, Yuanyuan Li, Zixu Wang, Yaoxing Chen, Yulan Dong

## Abstract

Pregnancy is a very complex and delicate process, inflammation in early pregnancy may result in pregnancy loss or defective implantation. Melatonin, mainly produced from the pineal body, which exerts several pharmacological effects. N6-methyladenosine (m6A) is the most prevalent modification of eukaryotic mRNA. The aim of this study was to investigate the association between melatonin and m6A during pregnancy and elaborate the underlying protective mechanism of melatonin during pregnancy. In vitro, melatonin was found to alleviated LPS-induced reductions in the number of implantation sites. Besides, melatonin was found to alleviate the activation of inflammation, autophagy and apoptosis pathways. In vitro studies demonstrated that melatonin regulated several downstream pathways in an m6A-dependent manner via melatonin receptor MTNR1B. Our findings revealed the important roles of m6A in the establishment of pregnancy and discovered a new mechanism of how melatonin protects pregnancy.

## Introduction

Compared to animals, humans have a lower success rate for pregnancy (about 30%), and 75% of pregnancy failures occur because of implantation failures [1]. Embryo implantation is an important step in the reproductive process. Implantation failure can be caused by many factors, such as maternal endocrine disorders, abnormal uterine development, inflammation and stress [2]. In addition to the immune system, many physiological systems are involved in coordinating pregnancy, including metabolic, endocrine, and circadian [3].

Melatonin, a neurohormone, is produced in the pineal, gut, skin, retina, ovary, placenta, glial cells, and lymphocytes (Tan et al., 2007). Melatonin has a wide range of regulatory and protective effects, such as synchronizing circadian rhythm [4], modulating the immune system [5], and regulating reproductive function [6]. In addition to its role in regulating circadian rhythms [7], researchers have demonstrated that melatonin present in high levels in human preovulatory follicular fluid [8], and regulate the timing of labor [9]. It has been revealed that melatonin enhanced embryo implantation by increases the E2 level during pregnancy [10], and melatonin receptor MT2 influence early gestation [11]. However, the mechanisms underlying the beneficial effects of melatonin on reproduction remain to be further elucidated.

N6-methyladenosine (m6A) is the most prevalent internal modification on mammalian RNA molecules [12]. The regulators of m6A include methyltransferases, demethylases and binding proteins, also known as writers, erasers and readers, respectively. Writers mainly include METTL3/14/16, WTAP and KIAA1429 [13], FTO and ALKBH5 are erasers [14]. The readers includes YTHDF1/2/3, YTHDC1/2, and IGF2BP1/2/3 preferentially recognize m6A on mRNA [15]. Emerging evidence implies that m6A methylation plays roles in many gynecological diseases, such as the adenomyosis [16] and endometriosis [17]. Our previous studies indicated that the m6A levels in the uterus increased as pregnancy progressed[18], that m6A methylation may be very important in the establishment of implantation and maintenance of pregnancy.

Although some studies have demonstrated that melatonin affects the female reproductive system, whether the protective effect of melatonin is exerted through the m6A methylation are still unknown. In the present study, we investigate the protective effects of melatonin on LPS-induced abnormal during early pregnancy. Our data demonstrate that melatonin protected mice from LPS-induced abnormal pregnancy, and alleviate inflammation, autophagy and apoptosis via MTNR1B-m6A.

## Results

### Melatonin protects against LPS-induced early pregnancy abnormalities in mice

Mice were treated as described in Figure 1A. The daily water consumption, feed intake and body weight of mice were monitored from D1 to D5 of gestation, the levels of daily water consumption (p<0.0001), feed intake (p<0.0001) and weight (p<0.0001) of mice was decreased immediately after LPS injection and melatonin alleviated the decrease (Figure 1B). Then we found the substantial decreased numbers of implantation sites in uterus from LPS group (p<0.0001), and the number of implantation sites and the wight of uterus in Mel+LPS group was higher than that in LPS group (p=0.0109) (Figure 1C-E). In addition, it was found that melatonin attenuated the LPS-induced decrease in blood glucose values (Figure 1F), as well as the decrease in platelet and white blood cell (Figure 1G). After that, melatonin was found to alleviate LPS-induced abnormal secretion of serum hormones, including estrogen (E2), progesterone (P4), corticosterone (CORT), and norepinephrine (NOR) (Figure 1H).

**Figure 1.**
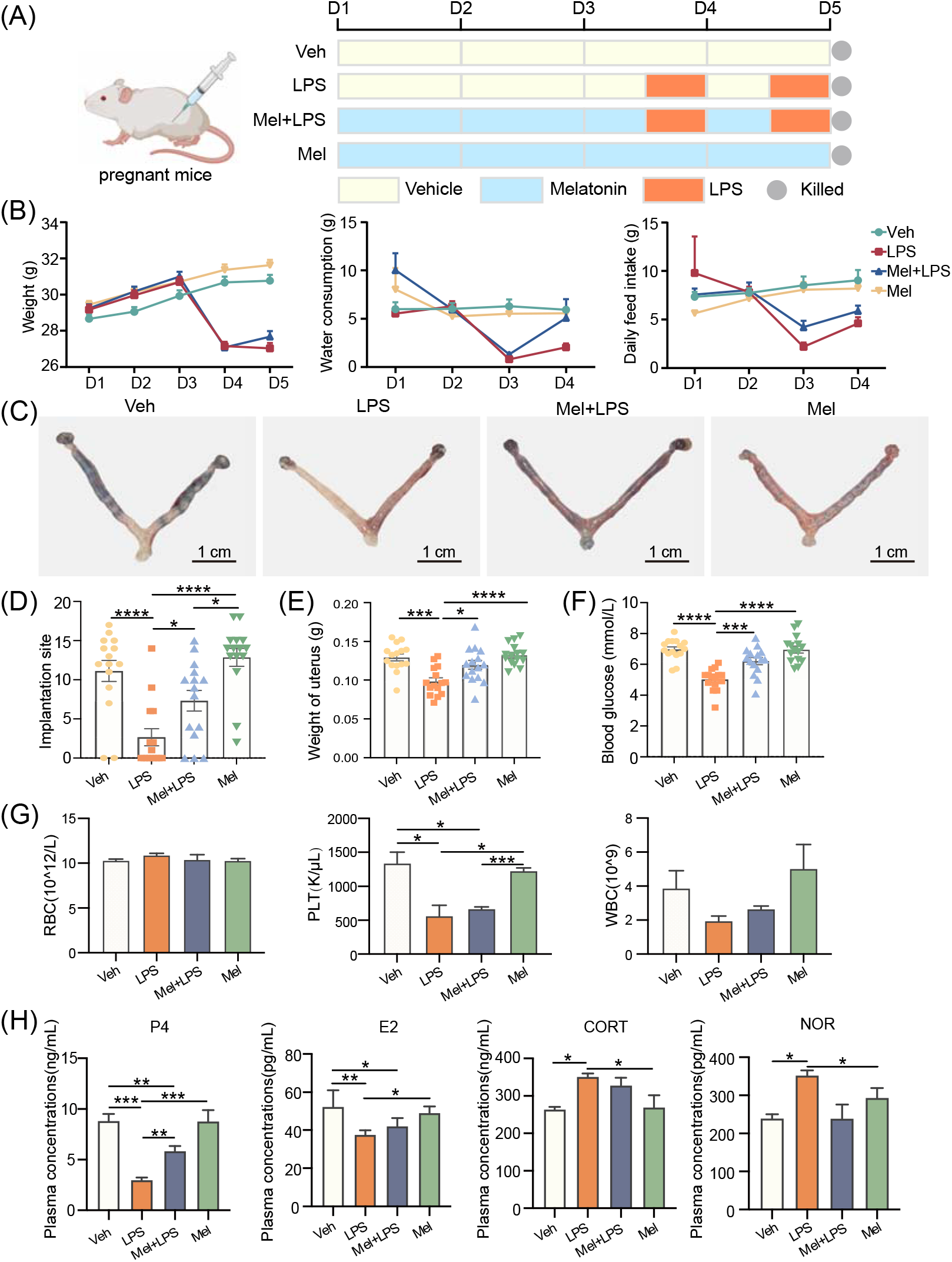
The protective effect of melatonin on mouse embryo implantation. (A) Schematic diagram of the animal experimental design. (B) Changes in the body weight, water cousumption and daily feed intake of pregnant mice. n=36 independent biological replicates. (C) Implantation sites was indicated by injecting Chicago Sky Blue dye. (D) The number of implantation sites in mice. n=15 independent biological replicates. (E) The weight of uterus in mice. n=15 independent biological replicates. (F) The blood glucose values in mice. n=15 independent biological replicates. (G) Blood was analyzed for the number of RBC, HB, PLT, WBC. RBC: red blood cell; HB: hemoglobin concentration; PLT: platelet count; white blood cell. n=3 independent biological replicates. (H) Serum hormone analysis in mice. n=3 independent biological replicates. P4: progesterone; E2: Estradiol-17β; CORT: Corticosterone; NOR: Noradrenaline. Veh: Vehicle treatment group; LPS: LPS treatment group; Mel+LPS: Melatonin and LPS co-treatment group; Mel: Melatonin treatment group; The data are presented as the mean ± SD. Levels of statistical significance for all data were determined by One-way ANOVA and Tukey test (* Indicates significant difference between the two groups; *p < 0.05; **p <0.01; ***p <0.001; ****p < 0.0001).

### Melatonin alleviates LPS-induced inflammation, autophagy and apoptosis in uterus

To explore the effect of LPS on gene expression in uterus, transcriptome sequencing data from LPS challenge and control mice were downloaded from the GEO database. We found 146 differentially expressed genes (DEGs) between the two groups (*Figure EV1*). The GO and KEGG analyses showed that these differentially expressed genes were mainly enriched in immunity and inflammation-related biological processes and pathways (Figure 2A and B). Furthermore, we examined the expression of 23 cytokines in uterus using a cytokine assay method. The results showed that melatonin significantly reduced the elevation of cytokines induced by LPS stimulation (Figure 2C and *Figure EV2*). Then, the similar patterns were observed in serum (*Figure EV3*). Subsequently, we analyzed inflammation, autophagy, and apoptosis related genes in decidua of mice in LPS challenge from the GEO database, these genes were found to be higher in the LPS-treated group (Figure 2D-F). In this study, the mRNA levels of several genes associated with inflammation, autophagy and apoptosis processes were detected (Figure 2G-I). The results showed that inflammation-related genes *Tlr4* (p=0.0432), *Myd88* (p=0.0220) and *Rela* (p=0.0052) were increased significantly in LPS group compare with the Veh group, while Mel+LPS has no significant effect compared to the Veh group. Additionally, melatonin alleviated the abnormal expression of autophagy and apoptosis genes after LPS stimulation, such as *Sqstm1*, *Becn1*, *Atg5*, *Atg7*, *Bcl2*, *Bax*, *Casp3*, *Casp9*.

**Figure 2.**
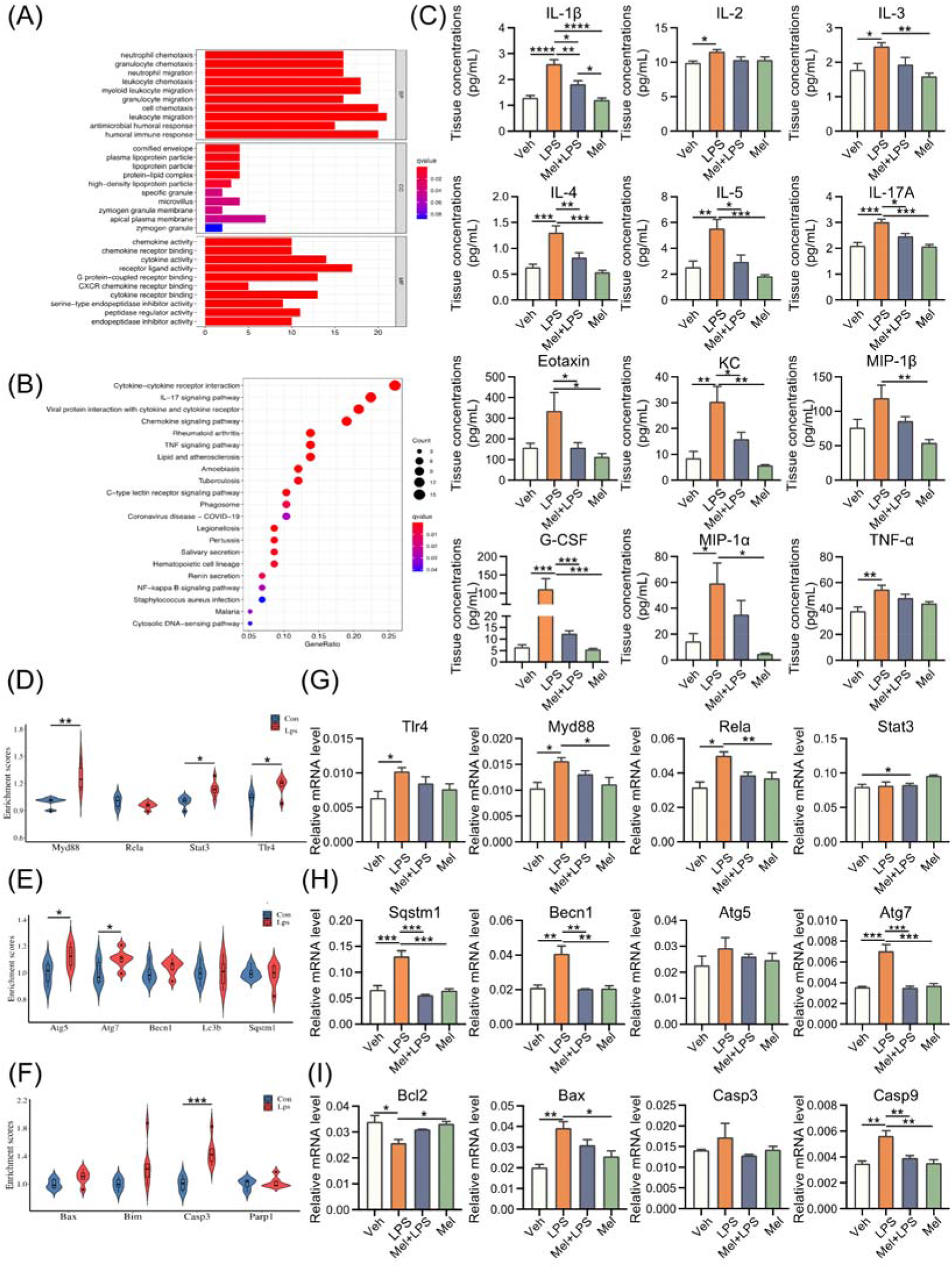
Melatonin alleviates LPS-induced inflammation, autophagy and apoptosis in uterus. (A) Gene Ontology (GO) enrichment analyses of DEGs between Con and Lps. GO terms were classified into three categories: biological process (BP), cellular component (CC), and molecular function (MF). (B) Kyoto Encyclopedia of Genes and Genomes (KEGG) pathway enrichment analyses of DEGs between Con and Lps. The top 21 pathways enriched in KEGG. (C) Uterus cytokine analysis by Luminex. n=4 independent biological replicates. (D) Violin plots show the expression levels inflammation-related genes mRNAs between Con and Lps. (E) Violin plots show the expression levels autophagy-related genes mRNAs between Con and Lps. n=3 independent biological replicates. (F) Violin plots show the expression levels apoptosis-related genes mRNAs between Con and Lps. n=3 independent biological replicates. (G) The mRNA levels of the inflammation-related genes in uterus of mice. n=3 independent biological replicates. (H) The mRNA levels of the autophagy-related genes in uterus of mice. n=3 independent biological replicates. (I) The mRNA levels of the apoptosis-related genes in uterus of mice. n=3 independent biological replicates. Veh: Vehicle treatment group; LPS: LPS treatment group; Mel+LPS: Melatonin and LPS co-treatment group; Mel: Melatonin treatment group; The data are presented as the mean ± SD. Levels of statistical significance for all data were determined by One-way ANOVA and Tukey test (* Indicates significant difference between the two groups; *p < 0.05; **p <0.01; ***p <0.001; ****p < 0.0001).

### Melatonin alleviates elevated m6A levels LPS-induced in uterus

To explore whether LPS affect m6A, we analyzed expression level of m6A regulators genes in decidua of mice in LPS challenge from the GEO database. Analysis result shown that the m6A writers *Mettl16* (p=0.04589) and the readers *Ythdc1* (p=0.0963), *Ythdf2* (p=0.0006) and *Ythdf3* (p<0.0001) were higher in the Lps group (Figure 3A). In our study, the m6A levels of the implantation sites were detected, the result showed melatonin alleviated the increase of m6A levels caused by LPS (Figure 3B), as well as the mRNA levels of m6A writers *Mettl3* and *Mettl14* (Figure 3C). Subsequently, we found that METTL3 expressed more strongly in LPS group (Figure 3D). Furthermore, the protein-protein connections were analyzed by STRING. The results suggested that melatonin receptor 1B (MTNR1B) was an interacting protein of the m6A recognition proteins (Figure 3E). In addition, the mRNA levels of *Mtnr1b* in the uterus were detected, the results showed that the expression of *Mtnr1b* tended to be enhanced after melatonin injection, suggesting that melatonin may function through its receptors (Figure 3F). The localization of METTL3, FTO, MTNR1B showed that they were specifically expressed in endometrial stromal cells (Figure 3G). Next, the m6A-seq was performed to detection on RNA from the uterus in the Veh, LPS and Mel+LPS groups. We found the differential m6A peaks were mainly enriched in the 3’UTR (Figure 4A and B). In addition, we analyzed the m6A motif that were changed after melatonin and LPS treatment (Figure 4C). Afterwards, differentially methylated genes between groups were showed (Figure 4D). Then we performed KEGG pathway analysis on the m6A-up genes in the LPS group, results showed pathways enriched in metabolic pathways and NF-κB signaling pathway (Figure 4E). In addition, the GO enrichment for m6A-up genes in the LPS group was performed (Figure 4F). Subsequently, the intersection of m6A-up genes in the LPS group and m6A-down genes in the Mel+LPS group was analyzed (Figure 4G). Then we analyzed the intersection of the m6A-down genes in the LPS group and the m6A-up genes in the Mel+LPS group (Figure 4H). Results showed that m6A modification of many genes was altered after melatonin treatment.

**Figure 3.**
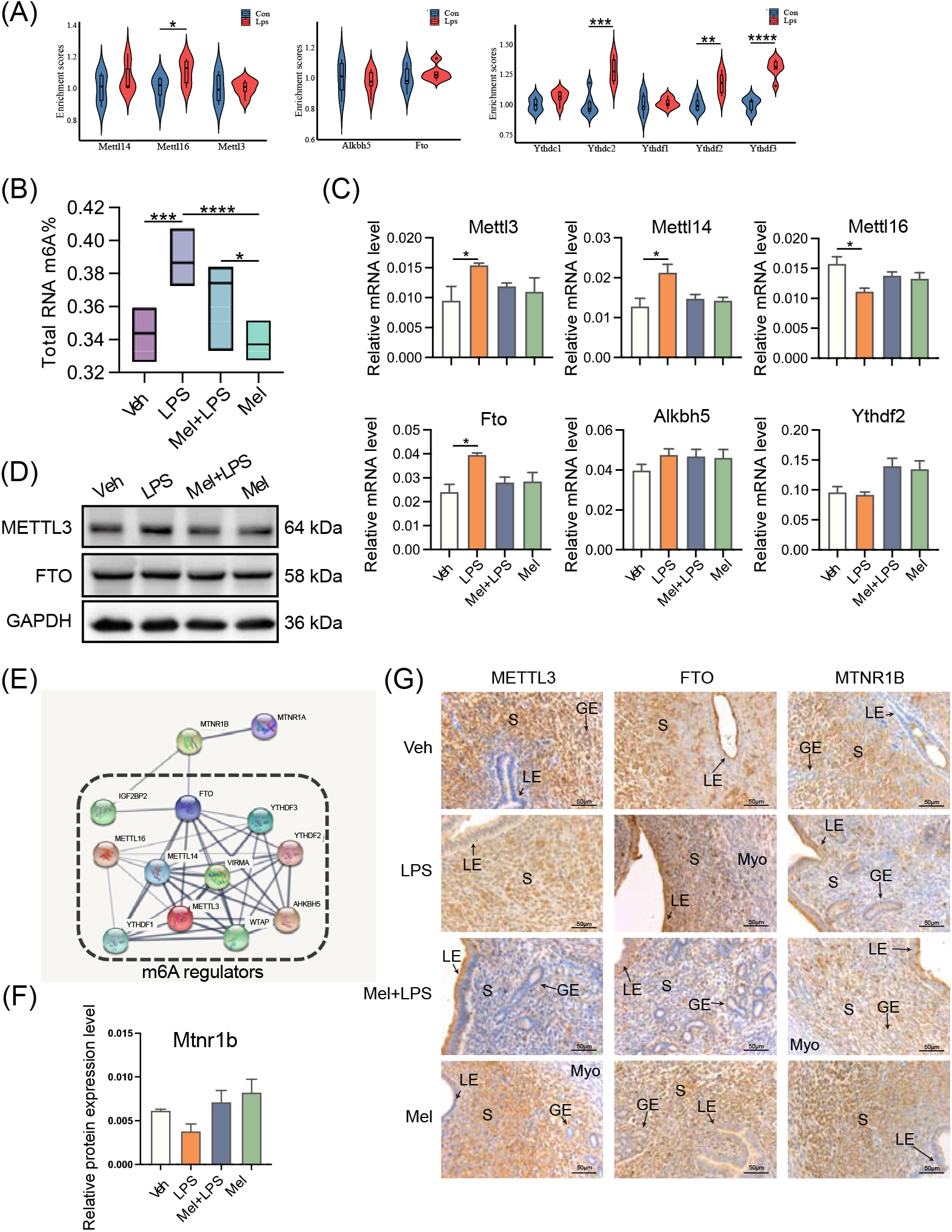
Melatonin alleviates LPS-induced elevated m6A levels in uterus. (A) Violin plots show the expression levels of m6A regulaters mRNAs between Con and Lps. n=3 independent biological replicates. (B) Global m6A levels in the uterus. n=6 independent biological replicates. (C) The mRNA levels of m6A regulaters in the uterus of mice. n=3 independent biological replicates. (D) Western blot bands of METTL3 and FTO. n=3 independent biological replicates. (E) Protein-protein interaction (PPI) network of the significant genes. Using the STRING online database. (F) The mRNA levels of *Mtnr1b* in the uterus of mice. n=3 independent biological replicates. (G) Immunohistochemical (IHC) staining of METTL3 and FTO in the uterus on D5. n=3 independent biological replicates. S: stroma; LE: luminal epithelium; GE: glandular epithelium; Myo: myometrium. Veh: Vehicle treatment group; LPS: LPS treatment group; Mel+LPS: Melatonin and LPS co-treatment group; Mel: Melatonin treatment group; The data are presented as the mean ± SD. Levels of statistical significance for all data were determined by One-way ANOVA and Tukey test (* Indicates significant difference between the two groups; *p < 0.05; **p <0.01; ***p <0.001; ****p < 0.0001).

**Figure 4.**
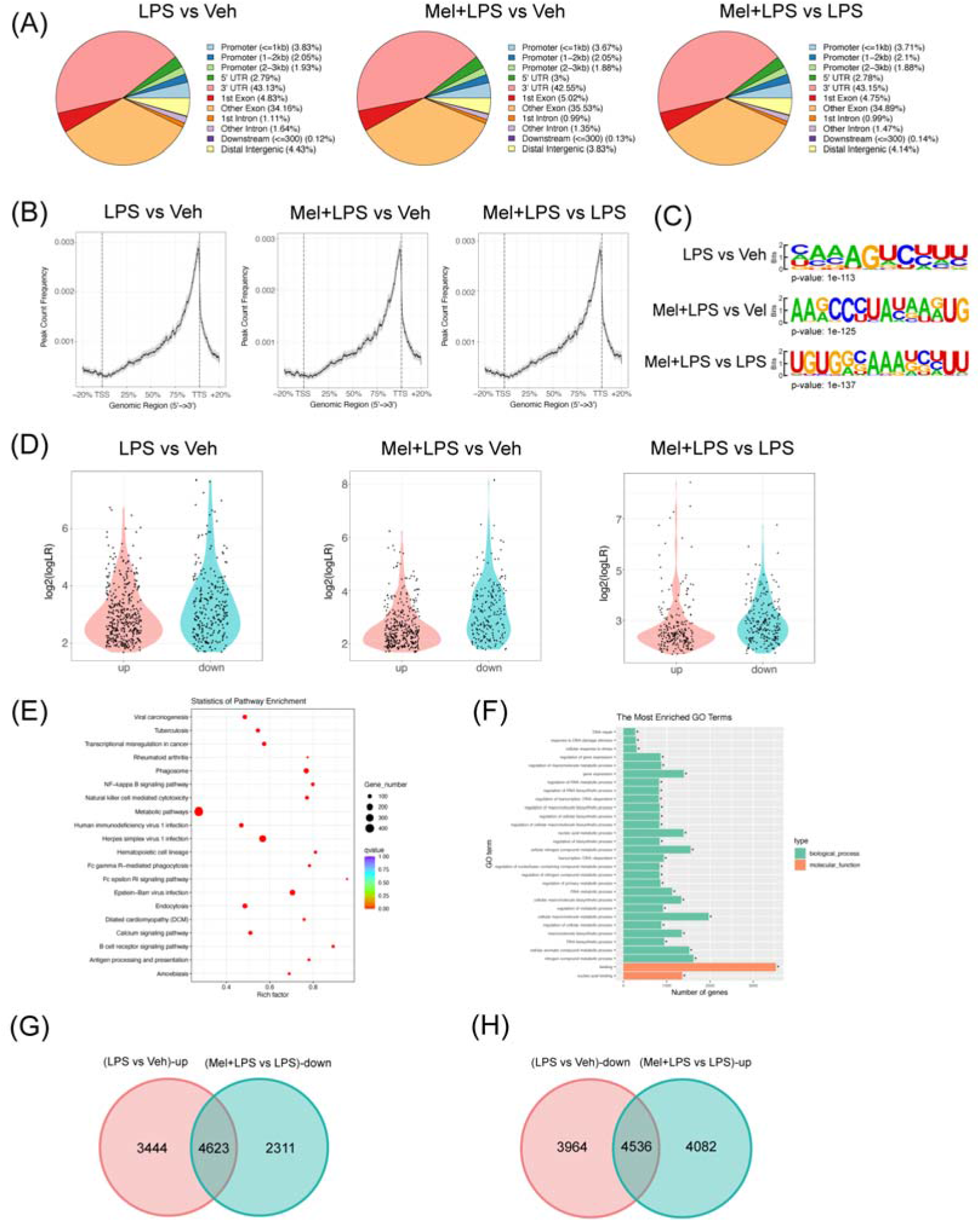
m6A-seq analysis of m6A modification after melatonin and LPS stimulation. (A) The annotation of m6A-seq different reads between different treatments. (B) Genome browser view of m6A-seq different reads between different treatments. (C) The motif of m6A-seq different reads between different treatments. (D) Violin plots of the m6A-up and m6A-down genes between different treatments. (E) KEGG pathway enrichment of m6A-up genes between LPS group and Veh group. (F) GO enrichment of m6A-up genes between LPS group and Veh group. (G) Venn diagram of m6A-up genes in the LPS group and m6A-down genes in the Mel+LPS group. (H) Venn diagram of m6A-down genes in the LPS group and m6A-up genes in the Mel+LPS group. n=3 independent biological replicates. Veh: Vehicle treatment group; LPS: LPS treatment group; Mel+LPS: Melatonin and LPS co-treatment group.

### Melatonin alleviates LPS-induced inflammation, autophagy and apoptosis in HESCs

Next, we use human endometrium cells (HESCs) to conduct in vitro experiments to study the mechanism of melatonin. CCK8 assay was performed to screen the valid concentrations of melatonin and LPS, melatonin (1 μM) and LPS (50 μg/mL) for 48 h was used in subsequent experiments (Figure 5A). Subsequently, we detected the protein levels of inflammation, autophagy and apoptosis expression of several genes associated with inflammation processes. Specifically, the inflammation related proteins p-RELA (p=0.0007) and ERK1/2 (p=0.0090), autophagy related proteins LC3B (p=0.0377) and ATG7 (p=0.0001), apoptosis related proteins c-PARP (p=0.0085), BAX (p=0.0080) and CASP1 (p=0.0188) were increased significantly in LPS group compared with the Veh group, while Mel+LPS has no significant effect compared to the Veh group (Figure 5B-D). Similarly, results of the transcript level test also showed that melatonin inhibited the activation of LPS on inflammation, autophagy, and apoptosis pathways (*Figure EV4A-C*). Overall, the in vitro test echoed the outcomes that were observed in the in vivo tests.

**Figure 5.**
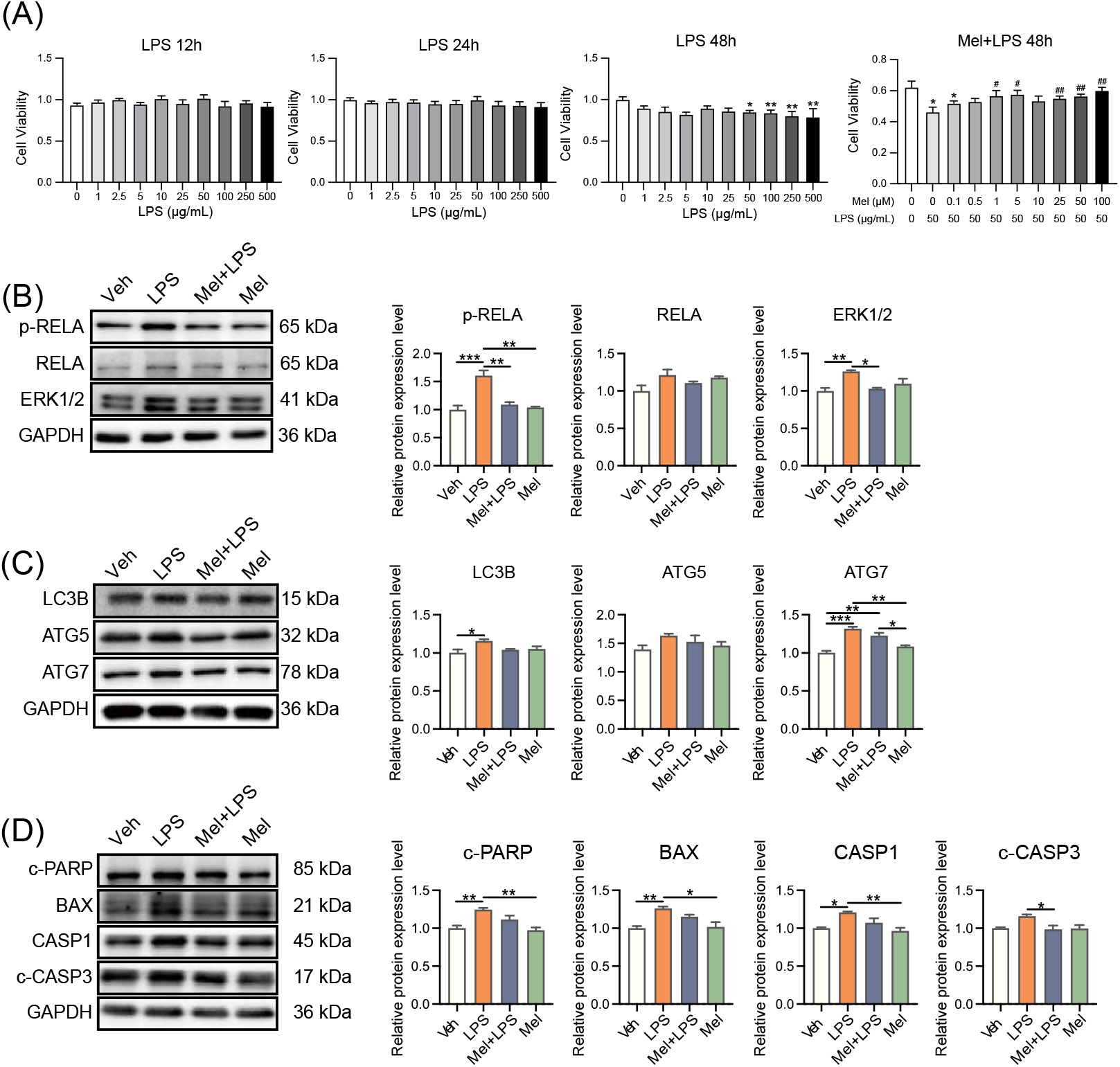
Melatonin alleviates LPS-induced inflammation, autophagy and apoptosis in human endometrial stromal cells. (A) Cell proliferation assay by CCK8 method. n=6 independent biological replicates. LPS: added LPS only; Mel+LPS: added melatonin and LPS; * Indicates a significant difference compared with the LPS 0 μg/mL group; *p < 0.05; **p < 0.01; ^#^ Indicates a significant difference compared with the LPS 50 μg/mL group; ^#^p < 0.05; ^##^p < 0.01. (B) Western blot bands of inflammation-related proteins in human endometrial stromal cells. n=3 independent biological replicates. (C) Western blot bands of autophagy -related proteins in human endometrial stromal cells. n=3 independent biological replicates. (D) Western blot bands of apoptosis -related proteins in human endometrial stromal cells. n=3 independent biological replicates. Veh: Vehicle treatment group; LPS: LPS treatment group; Mel+LPS: Melatonin and LPS co-treatment group; Mel: Melatonin treatment group; The data are presented as the mean ± SD. Levels of statistical significance for all data were determined by One-way ANOVA and Tukey test (* Indicates significant difference between the two groups; *p < 0.05; **p <0.01; ***p <0.001; ****p < 0.0001).

### Melatonin alleviates LPS-induced elevated m6A levels in HESCs

To explore the effects of LPS and melatonin on the m6A level of HESCs. Total RNA were extracted and m6A levels were detected by LC-MS/MS (Figure 6A), Result showed melatonin alleviated the m6A levels increase induced by LPS. As well as the mRNA levels of m6A regulators *METTL3*, *METTL14*, *ALKBH5*, *YTHDF1* (Figure 6B). Then the protein levels of m6A regulators were detected and found that METTL3 expressed more strongly in LPS group (*Figure EV4D*). Next, we detected the protein localization of METTL3 and FTO in HESCs by immunofluorescence and found that METTL3 was strongly expressed in the nucleus of HESCs regardless of the treatment conditions. Interestingly, the localization of FTO in the nucleus was reduced after LPS treatment, suggesting that melatonin may regulate m6A modification by controlling the entry of FTO into the nucleus (Figure 6C). we localized the protein of MTNR1B in HESCs and found that MTNR1B was localized in the nucleus, cytoplasm and cell membrane of HESCs (Figure 6D).

**Figure 6.**
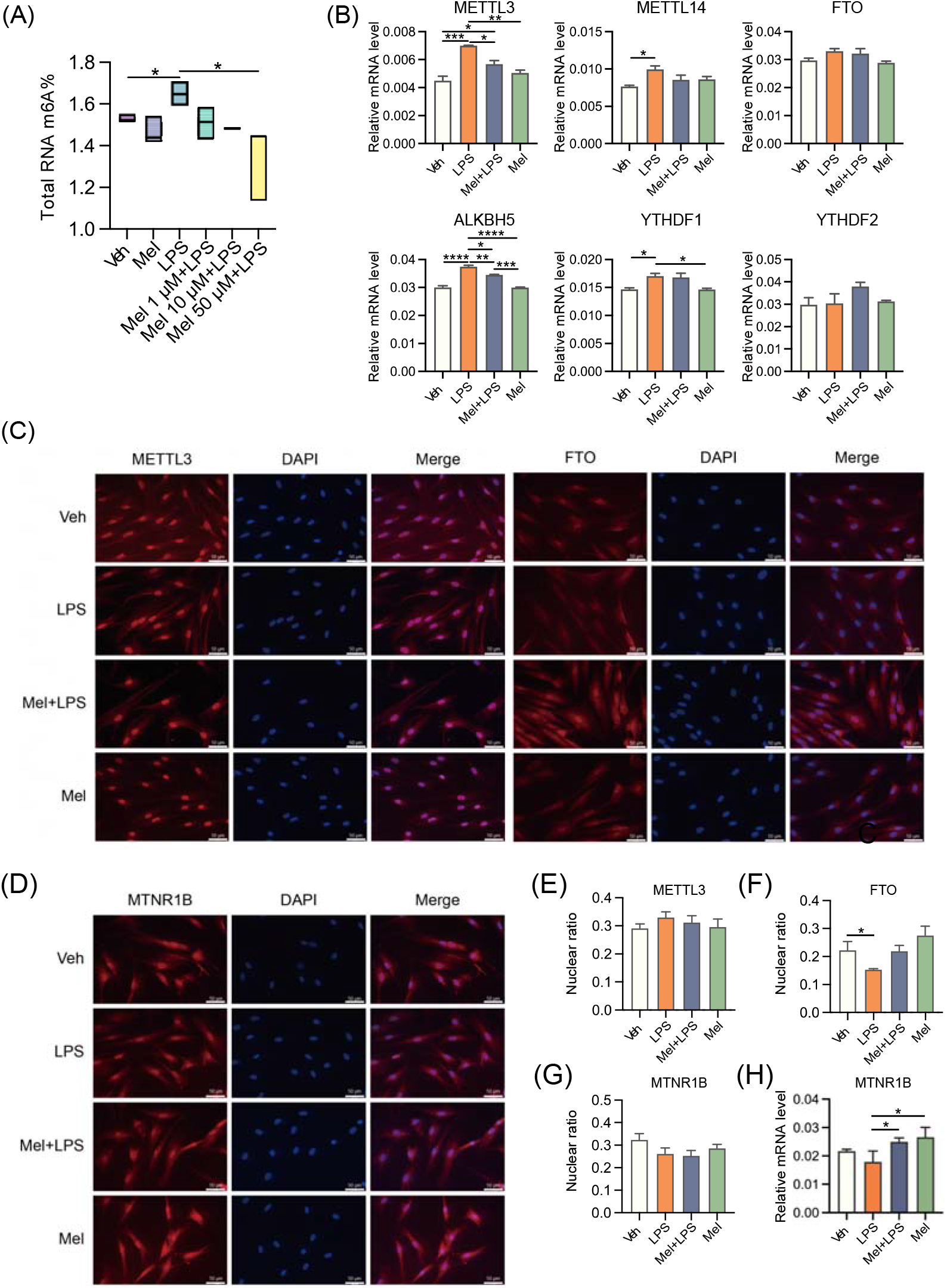
Melatonin alleviates LPS-induced elevated m6A levels in human endometrial stromal cells. (A) Global m6A levels of human endometrial stromal cells treated with LPS and different concentrations of melatonin. n=3 independent biological replicates. (B) The mRNA levels of the m6A regulators in uterus of human endometrial stromal cells treated with LPS and melatonin. n=3 independent biological replicates. (C) Immunofluorescence of m6A-related proteins in HESCs. (D) Immunofluorescence of MTNR1B in HESCs. (E) The ratio of the fluorescence intensity of METTL3 in the nucleus to the fluorescence intensity of METTL3 in the whole cell. n=6 independent biological replicates. (F) The ratio of the fluorescence intensity of FTO in the nucleus to the fluorescence intensity of FTO in the whole cell. n=6 independent biological replicates. (G) The ratio of the fluorescence intensity of MTNR1B in the nucleus to the fluorescence intensity of MTNR1B in the whole cell. n=6 independent biological replicates. (H) The mRNA levels of *MTNR1B* in the uterus of mice. The experiments were performed in triplicate, The data are presented as the mean ± SD. Levels of statistical significance for all data were determined by One-way ANOVA and Tukey test (* Indicates significant difference between the two groups; *p < 0.05; **p <0.01; ***p <0.001; ****p < 0.0001).

Then the ratio of fluorescence intensity in the nucleus was analyzed and found the localization of FTO in the nucleus decreased after LPS stimulation, while increased after melatonin addition, which may be related to its demethylation function (Figure 6E-F). It was also found that LPS and melatonin did not affect the nuclear expression of MTNR1B (Figure 6G). In addition, we examined the transcriptional level of MTRN1B in HESCs, the results showed the addition of melatonin activated the expression of MTNR1B (Figure 6H).

### Melatonin plays a protective role through MTNR1B

In order to explore whether melatonin protects cells through its receptor and m6A pathway, the melatonin receptor antagonist 4-P-PDOT and the METTL3-METTL14 complex inhibitor SAH were used. Cell viability was evaluated, 0.1 μM of 4-P-PDOT and 1 μM of SAH was used as the subsequent treatment concentrations (Figure 7A). Then the m6A levels were detected through LC-MS/MS, result showed that the ability of melatonin to regulate the level of m6A was weakened after antagonizing MTNR1B (Figure 7B). Subsequently, we found that inflammation, autophagy, and apoptosis related genes were increased significantly in LPS group compared with the vehicle group, while Mel+LPS group has no significant effect compared to the Veh group (Figure 7C-H). Additionally, the trend was confirmed by flow cytometric (Figure 7I).

**Figure 7.**
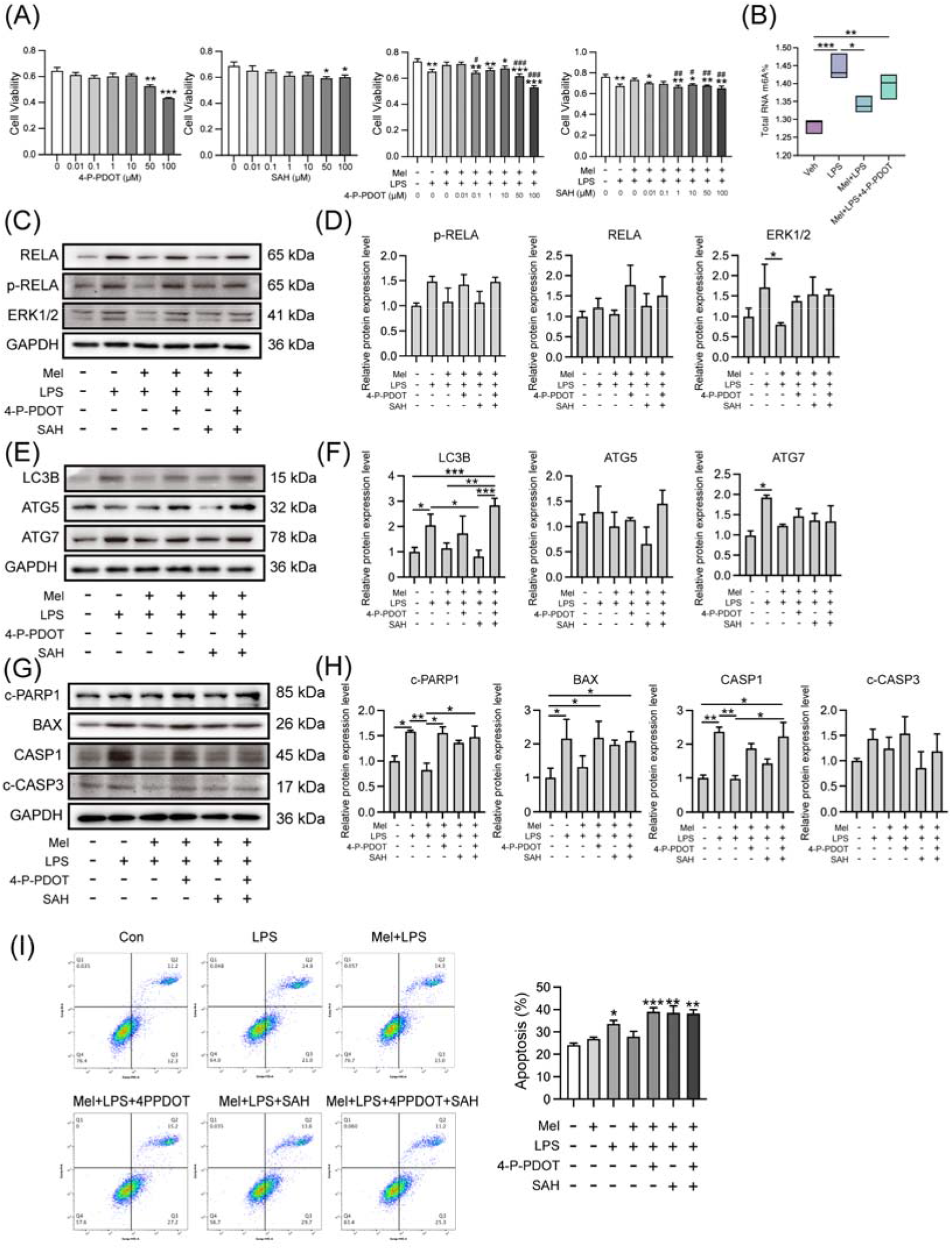
Melatonin plays a protective role through MTNR1B. (A) Cell proliferation assay by CCK8 method. n=6 independent biological replicates. 4-P-PDOT: 4-phenyl-2-propionamidotetralin, Melatonin receptor MTNR1B antagonist; SAH: S-adenosylhomocysteine, METTL3-METTL14 complex inhibitor; * Indicates a significant difference compared with the 4-P-PDOT/SAH 0 μM group; *p < 0.05; **p < 0.01; ***p < 0.001; ^#^ Indicates a significant difference compared with the LPS added group alone Difference; ^#^p < 0.05; ^##^p < 0.01; ^###^p < 0.001. (B) Global m6A levels of human endometrial stromal cells treated with 4PPDOT. n=3 independent biological replicates. (C) Western blot bands of inflammation-related proteins in human endometrial stromal cells. (D) Immunoblot analysis of p-RELA, RELA, ERK1/2. n=3 independent biological replicates. (E) Western blot bands of autophagy -related proteins in human endometrial stromal cells. n=3 independent biological replicates. (F) Immunoblot analysis of LC3B, ATG5, ATG7. n=3 independent biological replicates. (G) Western blot bands of apoptosis -related proteins in human endometrial stromal cells. n=3 independent biological replicates. (H) Immunoblot analysis of c-PARP1, BAX, CASP1, c-CASP3. n=3 independent biological replicates. (I) Flow cytometry was used to detect apoptosis after adding the 4PPDOT and melatonin, apoptosis cells were measured. n=3 independent biological replicates. Veh: Vehicle treatment group; LPS: LPS treatment group; Mel+LPS: Melatonin and LPS co-treatment group; Mel: Melatonin treatment group; The data are presented as the mean ± SD. Levels of statistical significance for all data were determined by One-way ANOVA and Tukey test (* Indicates significant difference between the two groups; *p < 0.05; **p <0.01; ***p <0.001; ****p < 0.0001).

## Discussion

Embryo implantation is an important step for the establishment of normal pregnancy, failed implantation is a major limiting factor in assisted reproduction [1]. There is now a lot of evidence for the beneficial effects of melatonin on pregnancy [19]. A previous study found that melatonin improve the implantation ability of mouse embryos, which may be related to the binding of uterine HB-EGF to ErbB1 and ErbB4 receptors [20]. Another experiment showed that melatonin improve embryo implantation and increase litter size by increasing E2 levels [10]. However, the mechanism of action of melatonin on inflammation in pregnancy remains to be supplemented.

In this study, LPS was used to induce abnormal pregnancy in mice during early pregnancy, then the protective effect of melatonin on pregnancy was investigated. The implantation of mice embryo was observed at D5, and melatonin was found to protect against the reduction in implantation sites induced by LPS stimulation. Since it has been reported that exogenous addition of melatonin down-regulate rat E2 levels and up-regulate P4 levels, it can also promote oocyte maturation in vitro and promote implantation of transplanted embryos [21]. In this study melatonin was found to alleviated the decrease in E2 and P4 and the increase in CORT and NOR induced by LPS. Earlier studies have demonstrated that LPS induce the inflammatory response and activate the NF-κB signaling pathway [22]. It has been reported that the anti-inflammatory mechanism of melatonin may include inhibition of TLR4 signaling [23]. In this study, melatonin was found to alleviated the activation of NF-κB signaling pathway caused by LPS. In other studies, the regulation of melatonin in autophagy has been observed in disease models [24]. Accordingly, in our study, autophagy pathway-related genes were detected, the results showed that melatonin decreased the autophagy pathway-related genes *Sqstm1*, *Becn1*, *Atg7* after LPS stimulation. It is suggested that melatonin play an active role by regulating autophagy. A large number of literatures have demonstrated the interaction between autophagy and apoptosis, and some studies have found that autophagy inhibit apoptosis [25]. Melatonin also prevents apoptosis by reducing CASP3 activation and reducing DNA damage[26]. In this study, apoptosis pathway-related genes were detected in the uterus of melatonin and LPS-treated mice. The results showed that melatonin could alleviate the abnormal increase of autophagy pathway-related genes Bax and Casp9 under LPS stimulation.

With the development of epigenetics, studies have shown that epigenetic regulation such as DNA methylation [27] and histone modification [28] are crucial for the female reproductive system, now RNA modification has become a new direction for studying reproductive function. A study found that the level of m6A in the endometrium of patients with adenomyosis was significantly reduced [16]. It has also been reported that m6A regulators are dysregulated in endometriosis [17]. Recent study has found that melatonin regulate the pluripotency of stem cells through the m6A pathway [29]. However, the mechanism of the effect of m6A on the pregnancy process has been rarely reported, and whether melatonin plays a role in improving pregnancy through the m6A pathway is still unknown.

In this study, protein interaction network analysis found that melatonin receptor MTNR1B interacts with m6A-related proteins, suggesting that melatonin may affect m6A modification through MTNR1B. The m6A level was detected, and found that the m6A level in the uterus increased after LPS stimulation, melatonin alleviated the abnormal increase. Then the transcription levels of m6A-related genes were detected, and it was found that melatonin alleviated the up-regulation of methylated genes *Mettl3* and *Mettl14*. The results of protein detection also indicated that the protein expression level of METTL3 was increased. In another study, it was also found that LPS stimulation resulted in up-regulation of m6A and METTL3 levels in human dental pulp cells [30]. The above results suggest that melatonin alleviate the LPS-induced increase of m6A levels in the uterus of mice by regulating m6A methylation-related genes. The in vivo results showed that intraperitoneal injection of melatonin in mice could alleviate LPS-induced abnormal pregnancy in mice, and melatonin could inhibit the abnormal increase of m6A modification, inflammation, autophagy and apoptosis in the uterus of mice.

Based on the above research results, this study selected human endometrial stromal cells for in vitro research and was committed to further exploring the specific mechanism of melatonin. The latest findings indicate that melatonin regulates stem cell pluripotency through the MT1-JAK2/STAT3-Zfp217 signaling axis [29]. In this study, the results showed that melatonin may affect m6A modification through its membrane receptor MTNR1B, then we selected MTNR1B antagonist 4-P-PDOT and METTL3-METTL14 complex inhibitor SAH for cell treatment. Then we found that the stabilizing effect of melatonin on m6A levels was inhibited when 4-P-PDOT was added, suggesting that melatonin may affect m6A through its receptor MTNR1B. In the present study, after treating cells with melatonin and LPS, it was found that melatonin could alleviate LPS-induced increases in the mRNA levels of inflammation-related genes *TLR4*, *MYD88*, *RELA*, *STAT3*, as well as the protein levels of RELA, p-RELA, ERK1/2 were increased. A similar situation was observed in a colitis study in rats, where melatonin reduced the TLR4, MyD88, and NF-κB up-regulation induced by trinitrobenzenesulfonic acid [31]. Interestingly, the alleviating effect of melatonin on LPS-induced inflammation was absent when MTNR1B was antagonized in the present study.

This study also conducted a series of tests on the autophagy pathway, and found that melatonin could alleviate the LPS-induced increase in the mRNA levels of autophagy-related genes *SQSTM1*, *LC3B*, *BECN1*, *ATG5*, *ATG7*, as well as the the protein levels of LC3B, ATG5, ATG7. However, when MTNR1B was antagonized, the alleviating effect of melatonin disappeared. In addition, we detected apoptosis of cells. The results showed that melatonin could alleviate the LPS-induced increase in the mRNA levels of apoptosis-related genes *BAX*, *BCL2*, *BIM*, *CASP3*, *CASP9*. as well as the the protein levels of c-PARP1, BCL2, CASP1, c-CASP3. The protective effect of melatonin on apoptosis was also inhibited after the addition of an antagonist of MTNR1B, which was confirmed by the flow cytometry. Interestingly, the expression of inflammation, autophagy, and apoptosis-related proteins was significantly reduced after the addition of SAH, possibly due to m6A being an important post-transcriptional modification that regulates mRNA translation [15]. When m6A methylation inhibited, these genes cannot be translated into proteins normally. In addition, it was found that the apoptosis of cells was significantly increased after the addition of SAH. Similarly, it has also been reported that inhibition of m6A modification lead to increased apoptosis. In previous studies on zebrafish embryos, it was found that METTL3 knockout results in embryonic tissue differentiation defects and increased apoptosis [32]. Studies have reported that silencing METTL3 in human osteosarcoma cells significantly inhibit cell proliferation, migration and invasion and promote apoptosis [33]. There are also reports of increased apoptosis following inhibition of the catalytic activity of METTL3-METTL14 [34]. The above results suggest that under normal circumstances, appropriate m6A modification maintain the physiological function of cells, while too high or too low m6A modification is not conducive to the survival of cells.

## Conclusions

This study found that melatonin alleviated LPS-induced implantation failure and abnormal pregnancy in mice through the MTNR1B-m6A pathway, which relieves the abnormal increase of inflammatory related proteins TLR4, MYD88, RELA, autophagy related proteins BECN1, ATG5 and ATG7, and apoptosis related proteins BAX, CASP9 and CASP3 (Figure 8).

**Figure 8.**
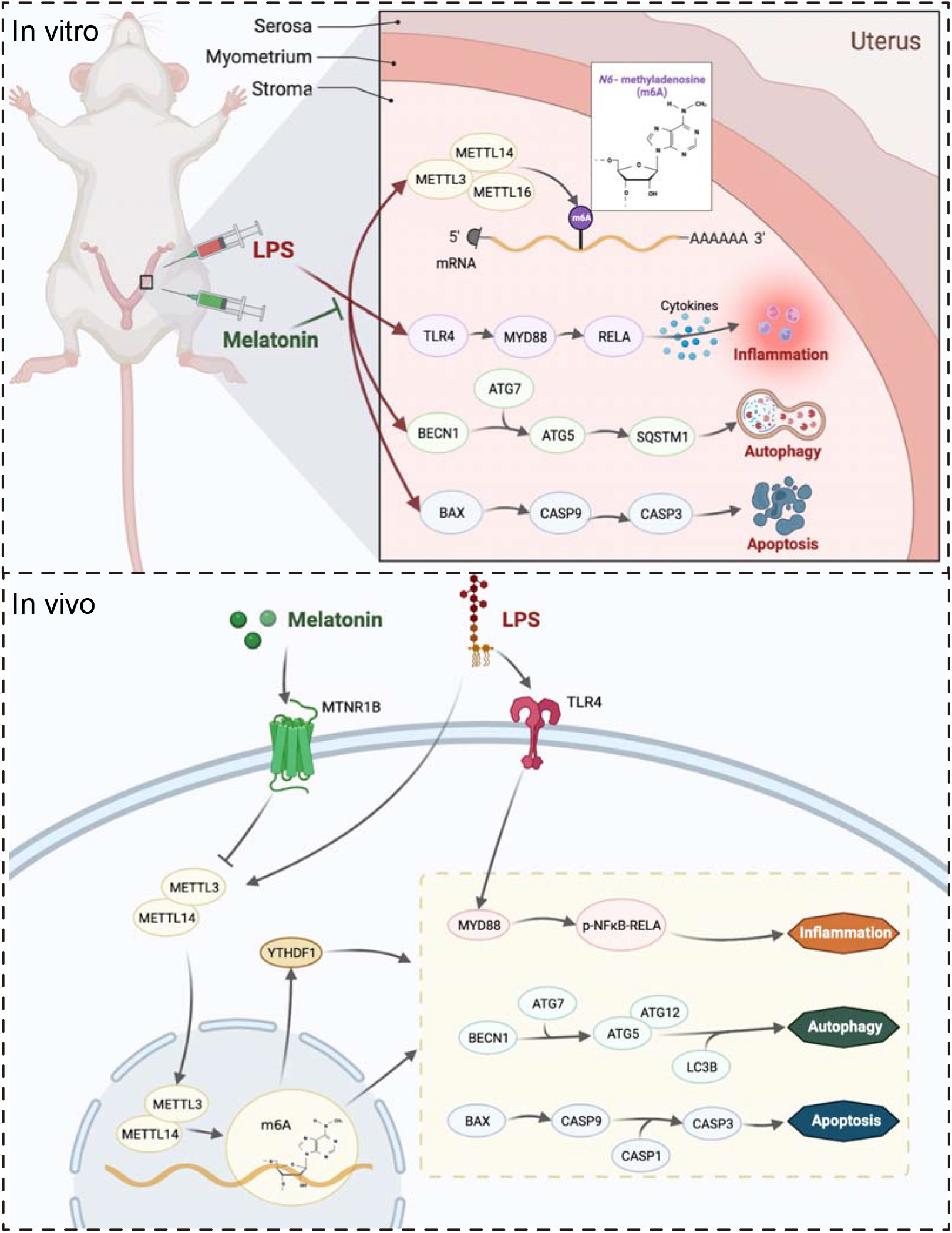
The mechanism by which melatonin protects pregnancy.

## Materials and Methods

### Animals and organ collection

Female ICR mice of 8 weeks old were purchased from Vital River Laboratory Animal Technology (Beijing, China). Mice were bred under controlled environmental conditions (12 h light/dark cycle, relative humidity 55-65% and 23-27°C). Female mice were mated to males at a ratio of 2:1. The day of mating was counted as day 0 (D0), and a vaginal plug was used to mark the first day of pregnancy (D1). As shown in Figure 1A, the pregnant mice were randomly divided into four groups: injected with vehicle (Veh), injected with LPS (LPS), injected with melatonin and LPS (Mel+LPS), injected with melatonin (Mel). The LPS (Sigma-Aldrich, WU, USA) injection volume was 0.5 mg/kg body weight dissolved in 0.9% sterile saline. The melatonin (Sigma-Aldrich, WU, USA) injection volume was 10mg/kg body weight dissolved in 0.9% sterile saline containing 3% ethanol. On D1 to D4, mice were received intraperitoneal injections of melatonin or vehicle daily at 9:00 am, and on days 3 and 4, LPS was injected intraperitoneally 30 minutes after the melatonin injection. The mice were sacrificed on D5, and the uterus was harvested. Blood samples were harvested, and blood routine examination was performed using the whole blood by a routine blood test instrument (Mindray, Shenzhen, China). Implantation sites were visualized after tail vein injection of 1% Chicago blue (Sigma-Aldrich, WU, USA).

### Ethics statement

All animal procedures were approved by the Institutional Animal Care and Use Committee (IACUC) of China Agricultural University (ethics approval number: AW03602202-2-1). All animal experiments followed the guidelines of the IACUC.

### Cell culture and treatment

Immortalized human endometrial stromal cell line (HESCs), CRL-4003, was the kind gift from Prof. Renwei Su (South China Agricultural University, China). Cells were cultured in DMEMB/F12 containing 10% FBS. Once cells were attached, HESCs were starved for 12 h in DMEM/F12, cells were treated with 1 μM Melatonin and 50 μg/mL LPS for 48 h. For experiments involving inhibitors, cells were incubated with inhibitor for 48 h. The following specific inhibitors were used: melatonin receptor antagonist 4-P-PDOT (MCE, NJ, USA) and the METTL3-METTL14 complex inhibitor SAH (MCE, NJ, USA). 4-P-PDOT was used at a concentration of 0.1 μM and SAH was used at a concentration of 1 μM.

### Blood index detection method

Routine blood examination indices were detected via a blood cell analyser (NIHON KOHDEN, Tokyo, Japan) in whole blood. The blood glucose of the ear vein was detected by a Yuyue blood glucose meter. The levels of progesterone (P4), Estradiol-17β (E2), Corticosterone (CORT), Noradrenaline (NOR) were measured spectrophotometrically in serum using radioimmunoassay kit according to the manufacturer’s instructions. These kits were purchased from Beijing Huaying Institute of Biotechnology.

### Luminex liquid suspension chip detection

The procedure was performed according to the instructions of Bio-Plex Pro Mouse Cytokine 23-plex (Bio-Rad, CA, USA). Serum was diluted 4 times with Sample Diluent, and tissue lysate samples were diluted 25 times with Sample Diluent. Finally, 50 μL of the diluted samples were loaded for detection, and then the standard was diluted according to the instructions. Dilute the microbeads with Assay Buffer. After the diluted microbeads are shaken, add 50 μL of each well to a 96-well plate, wash three times with a plate washer, and add 50 μL of the prepared standards, samples and Blanks to the 96-well plate., placed on a plate shaker protected from light, and incubated at room temperature for 30 min. Discard the sample, wash the plate 3 times, add 25 μL of diluted Detection Antibody to each well, place on a plate shaker to avoid light, and incubate at room temperature for 30 min. Discard the detection antibody, wash 3 times, use Assay Buffer to dilute Streptavidin-PE according to the instructions, add 50 μL of diluted Streptavidin-PE to each well, place on a plate shaker to avoid light, incubate at room temperature for 10 min, wash 3 times, each well. The wells were resuspended by adding 125 μL Assay Buffer, placed on a plate shaker at room temperature, shaken for 30 s in the dark, and detected by Bio-Plex MAGPIX System (Bio-Rad, CA, USA).

### Quantitative analysis of m6A level using LC-MS/MS

Total m6A was measured in 1 μg of total RNA extracted from the uterus using liquid chromatography-tandem mass spectrometry (LC-MS/MS). RNA was incubated with 1 μL of S1 nuclease (Takara, Okasa, Japan) for 4 h at 37°C. Then, 1 μL of alkaline phosphatase (Takara, Okasa, Japan) was added, and the reaction was incubated for 1 h at 37°C. The reaction mixture was extracted with chloroform. HPLC separation was performed using a C18 column (Shimadzu Corporation, Kyoto, Japan) with a flow rate of 0.2 mL/min at 35°C. Solvent A was 0.1% (vol/vol) formic acid in water, and solvent B was 0.1% (vol/vol) formic acid in methanol. A gradient of 5 min of 5% B, 10 min of 5-30% B, 5 min of 30-50% B, 3 min of 50-5% B, and 17 min of 5% B was used.

### m6A-seq

m6A-seq was performed as previously described (Dominissini et al., 2012). Briefly, 100 μg total RNA was extracted from the uterus using TRIzon Reagent (CWBIO, Jiangsu, China). mRNAs were fragmented into about 100-nt fragments and immunoprecipitated (IP) with 5 μg m6A antibody (Abcam, Cambridge, UK). The antibody-RNA complex was isolated by incubation with protein A beads (Invitrogen, CA, USA). All libraries were sequenced on Illumina Hiseq X10 (Illumina, CA, USA). Raw reads from Illumina sequencing were subjected to adaptor trimming and filtering of low-quality reads by Fastp (version 0.12.3). STAR (version 2.5.2a) was used to output a sorted genome-coordinate based Bam file. The exomePeak2 (version 4.2) was used to perform peak calling analysis on the IP and Input file. Difference peaks were analyzed using MACS2 (version 3.4). Peaks were annotated using CHIPseeker (version 1.22.1). Motif analysis was performed using HOMER (version 4.10). All sequencing data has been uploaded to Gene Expression Omnibus (GEO series record GSE216994).

### Identification and functional assessment of DEGs

Tissue-specific mRNA expression data was from public data sets (GSE152343). The EdgeR package (edgeR 3.14.0) and limma package (version 3.30.7) were used to identify differentially expressed genes (DEGs) between the LPS and control groups (p-value < 0.05 and fold change > 2). Heatmaps were generated using the pheatmap package (pheatmap version 0.7.7). Gene Ontology (GO) enrichment and Kyoto Encyclopedia of Genes and Genomes (KEGG) pathway analyses were performed with the packages “clusterProfiler”, “enrichplot” and “org.Hs.eg.db” (p-value < 0.05 and q-value < 0.05).

### Protein-protein interaction (PPI) network construction

The PPI network of identified DEGs was constructed using the STRING online database (http://string-db.org). Functional modules in interaction networks were identified using the Markov clustering algorithm. The most stringent protein interaction screening criteria were applied (confidence > 0.9).

### Immunohistochemistry

Paraffin-embedded tissue sections were used for examination of hematoxylin-eosin (HE) staining. For IHC staining, tissue sections were repaired in 0.1 M sodium citrate buffer. Slides were blocked with 3% hydrogen peroxide and 5% horse serum and then incubated with the following primary antibodies: anti-METTL3 (1:100, Proteintech, Wuhan, China), anti-FTO (1:100, Proteintech, Wuhan, China), anti-MTNR1B (1:100, Cell Signaling Technology, USA). Then, slides were incubated with secondary antibody. Chromogen detection was carried out with the DAB chromogen kit (Zhongshan Golden Bridge Biotechnology, Beijing, China). All images were taken using a microscope (Olympus, Tokyo, Japan).

### Immunofluorescence

Immunofluorescence staining was performed at room temperature. Cells were washed in PBS, then fixed with 4% paraformaldehyde for 30 min at room temperature and washed 3 times in PBS.Cells were permeabilized with 0.1% Triton-X100 (Sigma T8787) and washed 3 times in PBS. blocked in 5% goat serum in 37℃ for 30 min. Cells were blocked with 5% goat serum and then incubated with the following primary antibodies: anti-METTL3 (1:100, Proteintech, Wuhan, China), anti-FTO (1:100, Proteintech, Wuhan, China), anti-MTNR1B (1:100, abcam, Cambridge, UK). Then, cells were incubated with secondary antibody: goat anti-rabbit IgG H&L Alexa Fluor 488 (1:1000, cell signaling technology, MA, USA), for 1 hour in the dark. Coverslips were mounted to slides with DAPI. All images were taken using a microscope (Olympus, Tokyo, Japan).

### Total RNA extraction and quantitative real-time PCR (qPCR)

Total RNA was isolated with TRIzon Reagent (CWBIO, Jiangsu, China). Then, cDNA was synthesized with HiScript QRTsupermix for qPCR (+gDNA wiper) (Vazyme, Nanjing, Nanjing, China). Real-time quantitative PCR (qPCR) was performed with SYBR green master mix (Vazyme, Nanjing, China). Changes in fluorescence were monitored on a OneStep Plus instrument (Applied Biosystems, Waltham, USA). Relative gene expression was obtained by normalizing the expression results to *Gapdh* expression (The list of primers used are shown in *Supplementary Table1* and *Table2*).

**Table 1.**
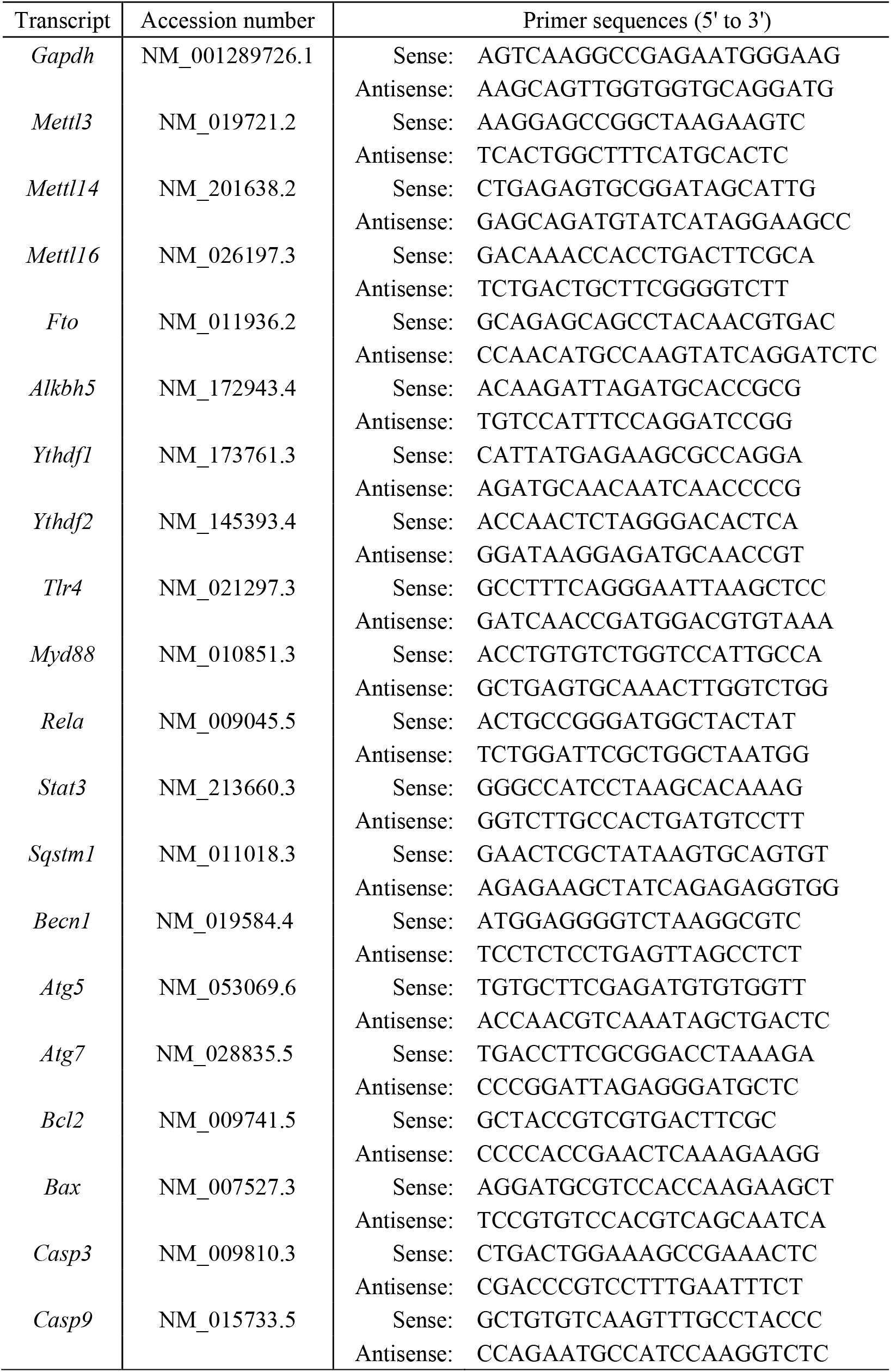
Primers for mouse used in this study.

**Table 2.**
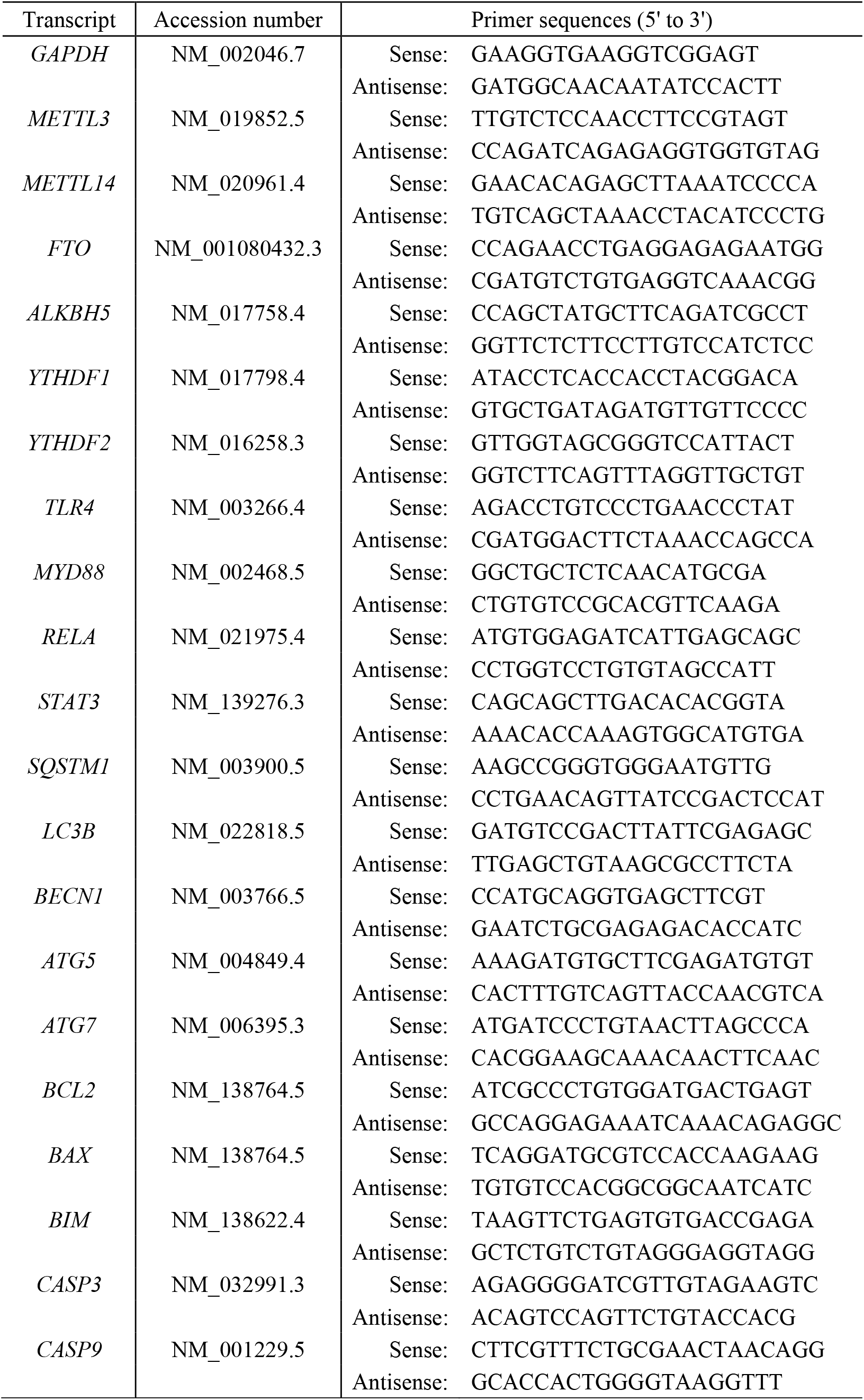
Primers for human used in this study.

### Western blot

Total protein was obtained from cells and tissues in RIPA lysis buffer (Biyuntian, Shanghai, China) with protease inhibitor (CWBIO, Jiangsu, China) and then separated by 10% SDS-PAGE. Membranes were blocked with 5% milk and then incubated with the following primary antibodies (for all antibodies, see *Supplementary Table3*), the concentrations were all 1:1000. Then the membranes were incubated with the secondary antibody anti-mouse or anti-rabbit horseradish peroxidase (HRP)-conjugated secondary antibodies (CWBIO, Jiangsu, China), the concentrations were 1:8000. Imaged with a Sapphire Biomolecular Imager (Azure Biosystems, CA, USA).

**Table 3.** Antibodies used in this study.

| Antibody | From |
| --- | --- |
| Rabbit anti-METTL3 polyclonal antibody | Proteintech |
| Rabbit anti-FTO polyclonal antibody | Proteintech |
| Rabbit anti-MTNR1B polyclonal antibody | Abcam |
| Mouse anti-GAPDH monoclonal antibody | Proteintech |
| Rabbit anti-RELA monoclonal antibody | Abclonal |
| Rabbit anti-phospho-RELA polyclonal antibody | Abclonal |
| Rabbit anti-ERK1/2 polyclonal antibody | Abmart |
| Rabbit anti-PARP1 polyclonal antibody | Cell Signaling Technology |
| Rabbit anti-BAX polyclonal antibody | Proteintech |
| Rabbit anti-Caspase1 polyclonal antibody | Proteintech |
| Rabbit anti-Cleaved-Caspase3 polyclonal antibody | Cell Signaling Technology |
| Rabbit anti-LC3B polyclonal antibody | Novus |
| Rabbit anti-ATG5 polyclonal antibody | Proteintech |
| Rabbit anti-ATG7 polyclonal antibody | Proteintech |

### CCK-8 cell viability assay

Cell counting kit 8 (CCK8) experiment for cell viability assay were performed using the CCK8 Kit (TargetMol, MA, USA). After a treatment, 10 μl of CCK8 solution was added to each well, and the cells were further incubated at 37 °C for 3 h. Absorbance was measured at 450nm.

### Flow cytometry to detect cell apoptosis

To quantify the cell apoptotic rate, cells were digested with 0.25% trypsin. Proteolysis was neutralized with 10% FBS, and the lysates were centrifuged at 3,000 rpm for 5 min, washed once with PBS, and stained using the Annexin V-FITC and a propidium iodide (PI) solution (Beyotime Biotechnology, China) for 15 min at room temperature away from light. The percentage of apoptotic cells for each sample was subsequently evaluated by a BD FACSCalibur flow cytometer (BD Biosciences, NJ, USA).

### Statistical analysis

Data analysis was performed by GraphPad Prism (version 8.1 for Windows, GraphPad) and R version 4.0.3 (R Development Team). Data are expressed as the means ± standard errors of the mean (SEMs). Violin plots were generated using the ggplot 2 package (version 2.2.1). p-values were calculated using Students t-test or analysis of variance (ANOVA) with Dunnetts test (for one-way ANOVA) or Tukeys (for two-way ANOVA) multiple-comparison test, with statistical significance as follows: *p<0.05; **p<0.01; ***p<0.001, ****p<0.001.

## Author contributions

**Shisu Zhao:** Data curation, Investigation, Methodology, Visualization, Writing - original draft, Writing - review & editing. **Yanjun Dong:** Investigation, Writing - review & editing. **Yuanyuan Li:** Software, Investigation. **Yaoxing Chen:** Investigation, Writing - review & editing. **Zixu Wang:** Investigation, Writing - review & editing. **Yulan Dong:** Funding acquisition, Investigation, Methodology, Writing - review & editing.

## Conflict of interest

The authors declare that they have no known competing financial interests or personal relationships that could have appeared to influence the work reported in this paper.

## Acknowledgments

We thank Prof. Renwei Su (South China Agricultural University, China) for providing CRL-4003 cells. The present work was supported by the National Natural Science Foundation of China (grant nos. 31972633, 31572476 and 31272483) and National Natural Science Foundation of Beijing (grant nos. 6172022 and 6212018).

## The paper explained

### Problem

A large number of women around the world suffer from abnormal pregnancies. Melatonin, as an anti-inflammatory hormone, has a protective effect on mammalian pregnancy, but its mechanism remains unclear. N6-methyladenosine (m6A) is the most prevalent internal modification on mammalian RNA molecules. Whether melatonin works through the m6A pathway remains to be investigated.

### Results

Melatonin alleviated LPS-induced implantation failure and abnormal pregnancy in mice through the MTNR1B-m6A pathway, relieves the increase expression of inflammatory, autophagy, apoptosis related proteins.

### Impact

This study shows the prospect of using melatonin as a pregnancy protectant. In addition, this study clarified that m6A modification is involved in the regulation of pregnancy process, which may be considered as a new therapeutic target for infertility.

## Data availability

The m6A-seq raw sequence data has been uploaded to Gene Expression Omnibus GSE216994 (https://www.ncbi.nlm.nih.gov/geo/query/acc.cgi?acc=GSE216994).

---

**Figure EV1.**
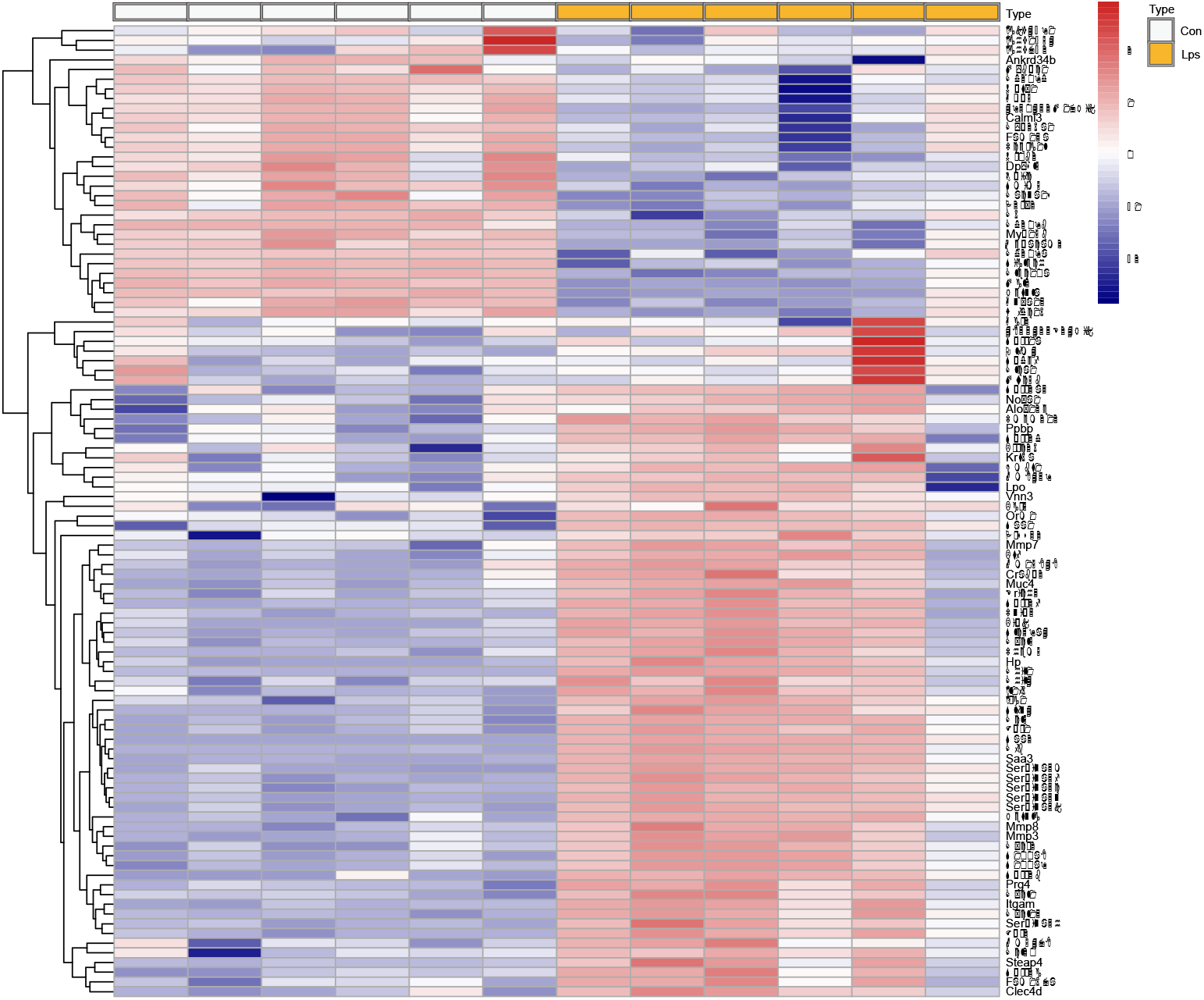
The differentially expression genes in decidua of mice in LPS challenge. Heatmap of differentially expressed genes (DEGs) in GEO datasets. The heatmap shows the top 100 genes with the most significant DEGs between the endometrium of the Con and LPS (Con: control group, Lps: LPS challenge group). n=3 independent biological replicates.

**Figure EV2.**
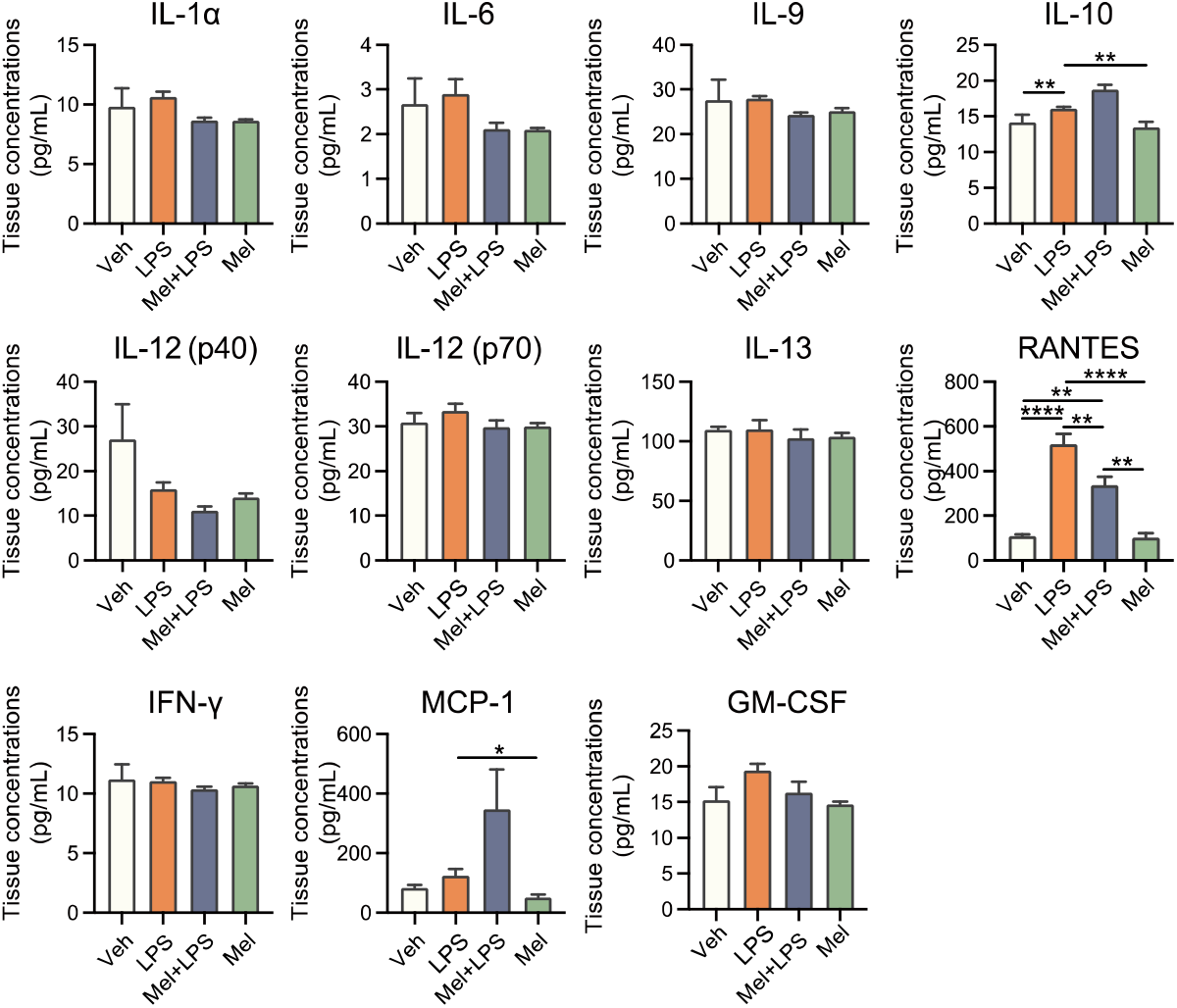
Mice uterus cytokine analysis by Luminex. n=4 independent biological replicates. Veh: Vehicle treatment group; LPS: LPS treatment group; Mel+LPS: Melatonin and LPS co-treatment group; Mel: Melatonin treatment group; The data are presented as the mean ± SD. Levels of statistical significance for all data were determined by One-way ANOVA and Tukey test (* Indicates significant difference between the two groups; *p < 0.05; **p <0.01; ***p <0.001; ****p < 0.0001).

**Figure EV3.**
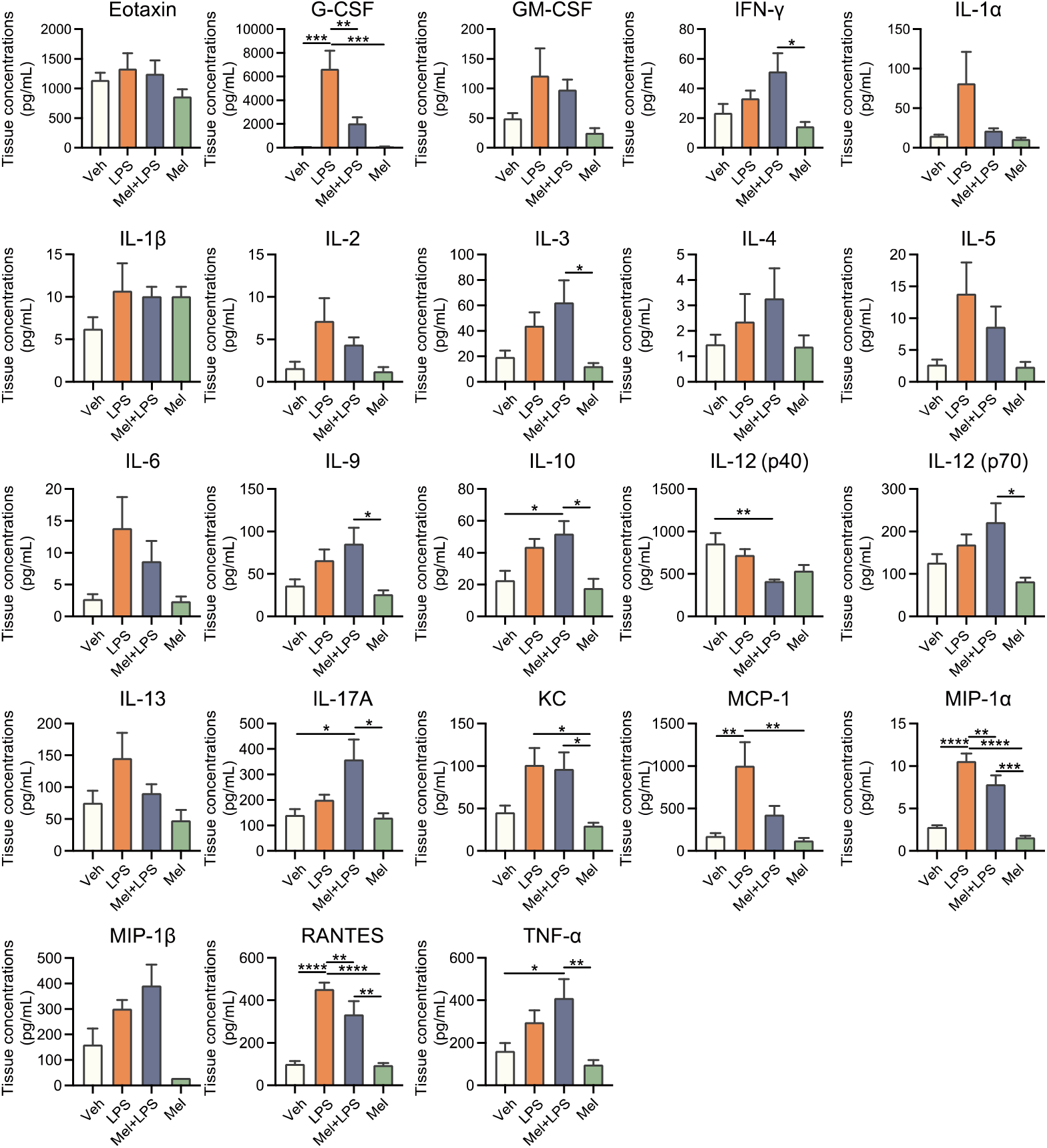
Serum cytokine analysis by Luminex. n=4 independent biological replicates. Veh: Vehicle treatment group; LPS: LPS treatment group; Mel+LPS: Melatonin and LPS co-treatment group; Mel: Melatonin treatment group; The data are presented as the mean ± SD. Levels of statistical significance for all data were determined by One-way ANOVA and Tukey test (* Indicates significant difference between the two groups; *p < 0.05; **p <0.01; ***p <0.001; ****p < 0.0001).

**Figure EV4.**
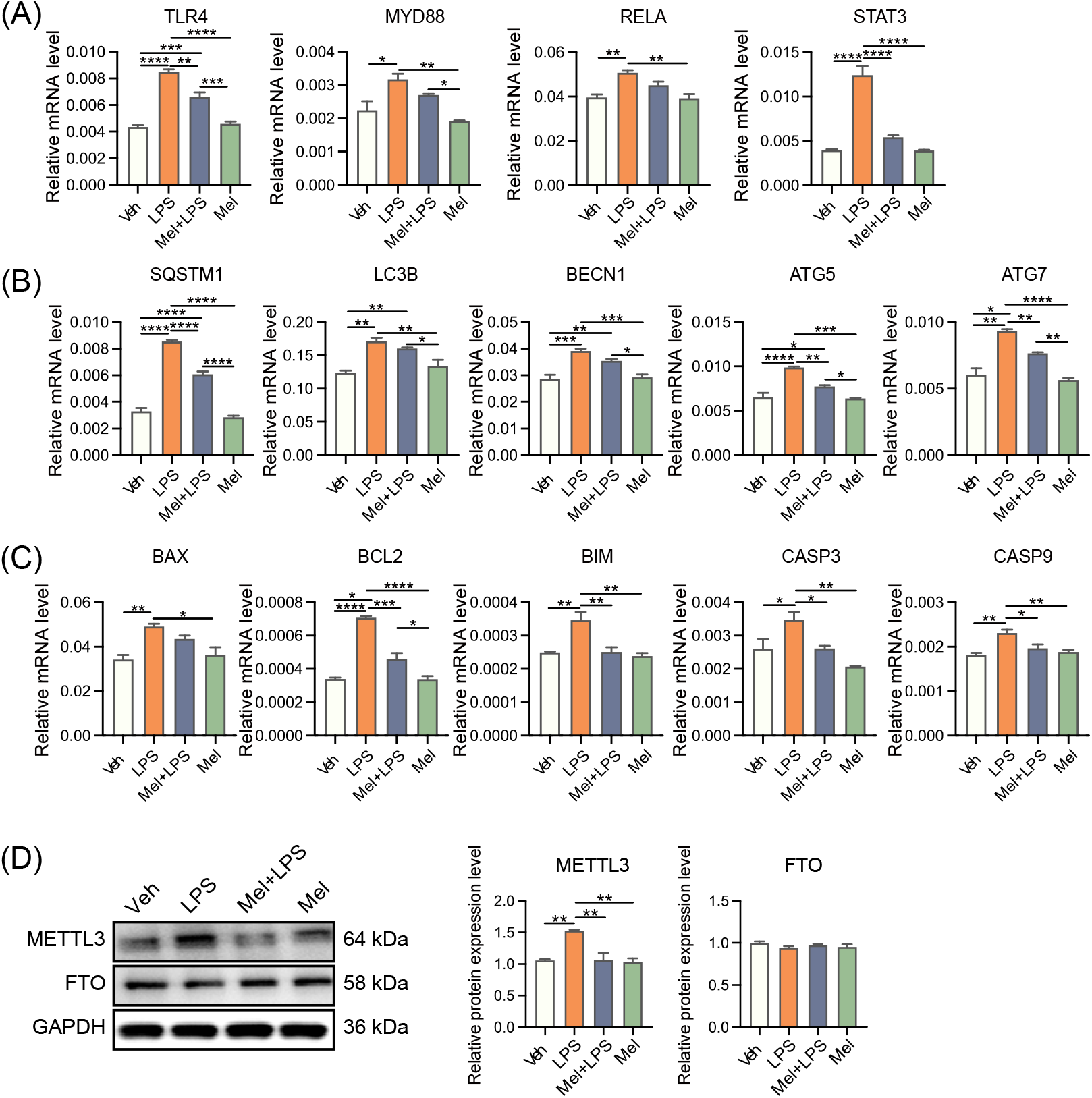
Transcript and protein levels detection in HESCs treated with melatonin and LPS. (A) The mRNA levels of the inflammation-related genes in human endometrial stromal cells. n=3 independent biological replicates. (B) The mRNA levels of the autophagy-related genes in human endometrial stromal cells. n=3 independent biological replicates. (C) The mRNA levels of the apoptosis-related genes in human endometrial stromal cells. n=3 independent biological replicates. (D) Western blot bands of METTL3 and FTO in human endometrial stromal cells treated with LPS and melatonin. n=3 independent biological replicates. Veh: Vehicle treatment group; LPS: LPS treatment group; Mel+LPS: Melatonin and LPS co-treatment group; Mel: Melatonin treatment group; The data are presented as the mean ± SD. Levels of statistical significance for all data were determined by One-way ANOVA and Tukey test (* Indicates significant difference between the two groups; *p < 0.05; **p <0.01; ***p <0.001; ****p < 0.0001).

## Notes

### Competing Interest Statement

The authors have declared no competing interest.

